# Direct Evidence of Effect of Glycerol on Hydration and Helix-to-Sheet Transition of Myoglobin

**DOI:** 10.1101/275321

**Authors:** M. Hirai, S. Ajito, M. Sugiyama, H. Iwase, S.-I. Takata, N. Shimizu, N. Igarashi, A. Martel, L. Porcar

## Abstract

By using wide-angle X-ray scattering (WAXS), small-angle neutron scattering, and theoretical scattering function simulation, we have clarified the effect of glycerol on both the thermal structure transition and the hydration-shell of myoglobin. At the glycerol concentration, ≤ ∼40 % v/v, the decreasing tendency in the maximum dimension and the radius of gyration was observed by X-ray scattering. The neutron scattering result using the inverse contrast variation method directly shows the preservation of the hydration-shell density at the concentration ≤ ∼40 % v/v. This phenomenon is reasonably explained by the preferential exclusion of glycerol from the protein surface to preserve the hydration shell, as suggested by the previous studies. While, at the concentration, ≥ 50 % v/v, the opposite tendency was observed. It suggests the preferential solvation (partial preferential penetration or replacement of glycerol into or with hydration-shell water surrounding the protein surface) occurs at the higher concentration. The observed WAXS scattering data covers the distinct hierarchical structural levels of myoglobin structure ranging from the tertiary structure to the secondary one. Therefore, we have clarified the effect of glycerol on the thermal structural stability myoglobin at different hierarchical structural levels separately. Against the temperature rise, the structural transition temperatures for all hierarchical structural levels were elevated. Especially, the tertiary structure of myoglobin was more stabilized compared with the internal-structure and the helix-to-cross transition. It suggests that the protective action of glycerol on protein structures essentially results from the preservation of the preferential hydration-shell of proteins.

## 1. Introduction

As is well-known, glycerol is a simple polyol compound that is widely used as a nontoxic additive in various industrial products for preservative, humectant, and thickening stabilizer, and also used as a cryoprotectant for storing enzymatic reagents, bacteria, nematodes, mammalian embryos and so on. The studies on the function of protein stabilization by glycerol have been conducted since long ago. The early study of the effect of glycerol on proteins using the densitometric measurement reported that the preferential hydration of proteins in glycerol-water mixture minimizes the surface of contact between proteins and glycerol to stabilize the native structure of globular proteins [1]. The H-exhange exchange study of myoglobin indicated that the protein fluctuation is hardly affected by solvent glycerol [2]. On the other hand, the static light scattering study suggested that the repulsive intermolecular force increase with glycerol concentration, which is explained by the incorporation of a layer of structured water that is enhanced in the presence of glycerol [3]. The molecular dynamics simulation suggested that preferential hydration of proteins in glycerol/water mixtures mainly originates from electrostatic interactions that induce orientations of glycerol molecules at the protein surface such as that glycerol is further excluded [4]. This study also suggested that glycerol prevented protein aggregation by inhibiting protein unfolding and by stabilizing aggregation-prone intermediates through preferential interactions with hydrophobic surface regions that favor amphiphilic interface orientations of glycerol. The Ramman and quasi-elastic inelastic neutron scattering study of the effect of glycerol on lysozyme showed that the protein’s fast conformational fluctuations and low-frequency vibrations and its temperature variations are very sensitive to behavior of the solvents [5]. The recent rheometer study on the effect of glycerol on the dynamics of collagen re(de)naturation suggests the presence of a nanometer thick, glycerol-free hydration layer where glycerol is fully expelled out of it [6]. In spite of many previous studies, there are fewer direct evidence showing the effect of glycerol on the protein structure, its preferential hydration layer and thermal stability.

On the other hand, the stabilities of proteins in aqueous solutions have been shown to vary by the addition of salts and neutral substances and also by the changes in temperature and pressure [7, 8]. In other words, structures and functions of proteins are controlled by a marginal balance between the stabilizing factor and the destabilizing one. Such a marginal balance would be possible disturbed or changed by external environmental factors or additives. Especially, since the interior of living cells is in so-called molecular-crowding environments where amounts of various types of small and large molecules coexist [9], the effect of the molecular-crowding environment on the structures and functions of proteins has been focused recently. The molecular-crowding environment would evidently affect the equilibrium states of proteins from the physicochemical point of view, whereas, interpretations of molecular-crowding effect remain controversial [10]. In addition, hydration of biopolymers is basically determined by characteristics of low molecules constituting them, and it is also considered to be the key determinant for isothermal, concentration-dependent effects on protein equilibria [11-13]. As shown by inelastic neutron scattering [14-18], the dynamics of proteins is coupled with and/or governed by water molecules surrounding them. Therefore, it is essentially important to clarify the relation between the protein hydration and the molecular-crowding environment even for small molecules, not only theoretically but also experimentally.

From the physicochemical interpretations of the small-molecular-crowding effect on a protein [10-12], a change in the chemical potential of bulk water due to a highly concentrated crowder-molecules causes a difference in chemical potential with water at the protein interface, and osmotic stress are generated. In order to relax the distortion by the osmotic stress, the surface structure and the exclusion area (preferential hydration area) of the protein change to shift to another equilibrium state different from that in the dilute solution, and at the same time, the changes in the diffusive motion (translational/rotational motion, internal motion) and in the intermolecular interaction are also induced. Thus, clearly the small-molecular-crowding environments cause changes of the equilibrium states of proteins [10, 19]. However, such a physicochemical interpretation does not still specify what kind of structural state a protein will be realized. In addition, the experimental observation of the structure of a protein in a solution where amounts of other molecules coexist is generally difficult because the significant increase of background scattering from co-solutes disturbs the observation of high-statistical scattering data of a protein. Therefore, most of the experimental studies of protein structure and stability were conducted under dilute-solution conditions [7, 8], and structure measurements of proteins under molecular-crowding environments have been rarely done. In contrast, although the remarkable progress of molecular-dynamics simulations in recent years has served fruitful insights on the effect of molecular-crowding on protein structure and dynamics [20-23], enough evidence of the changes of the structural stability and hydration of proteins caused by molecular-crowding has not been shown yet.

In the present study, we have clarified the effect of glycerol on both the hydration-shell and thermal structure transition of myoglobin by using the synchrotron radiation wide-angle X-ray scattering (SR-WAXS) method and the small-angle neutron scattering (SANS) method. Myoglobin is one of the good examples in the studies of protein folding[7] since it shows the cross-beta transition accompanying amyloid aggregate formation [24, 25]. As shown previously, the SR-WAXS method enables us to observe the whole hierarchical structure of proteins from their quaternary or tertiary structures to secondary one in solutions [26] and to analyze the details of the structural transition process of proteins separately at each hierarchical structure level [27]. We have already clarified the initial process of amyloid formation of apomyoglobin by using the SR-SAXS [28], and the characteristics of dynamics of apomyoglobin in the helix-to-sheet transition by the combination of SR-WAXS method and the elastic incoherent neutron scattering method [29]. In the neutron scattering measurements, we have employed the inverse-contrast variation method that is quite unique to elucidate selectively only the structures of the biological materials we focus [30, 31].

We have succeeded to obtain scattering data of myoglobin with high statistical accuracy in concentrated glycerol solutions (∼75 % v/v). By the combination of experimental results and theoretical solution scattering simulations, we have obtained a direct structural evidence on the effect of glycerol on the hydration-shell of myoglobin and the initial process of amyloid transition of myoglobin induced by temperature variation. The present results clearly indicate that the function of glycerol as a stabilizer of proteins is related to the preservation of protein hydration by the preferential exclusion of glycerol from the hydration-shell region of the protein at low glycerol-concentration (≤ ∼40 % v/v), which agree well with the previous studies using different techniques [1-6]. The results at high glycerol-concentration (≥ ∼50 % v/v), suggests the appearance of the preferential solvation (partial preferential penetration or replacement of glycerol into or with hydration-shell water surrounding the protein surface).

## 2. Experimental

Myoglobin from horse skeletal muscle, glycerol and deuterated glycerol (98 atom % D) were purchased from SIGMA Chemical Co. (USA) and used without further purification. All other chemicals used were of analytical grade. The deuterium oxide (99.9 atom % D, SIGMA) was used for neutron scattering experiments. The buffer solvent used was 10 mM HEPES (*N*-(2-hydroxymethyl) piperazine-*N*′-(2-ethane-sulfonic acid)) at pH 7.4. The myoglobin was dissolved in the buffer solvent on the concentration of 5 % w/v, which was used as the protein stock solution. The glycerol solutions with different concentrations were prepared. The protein solution and the glycerol solution were mixed by appropriate ratios. Finally, we obtained the protein solutions (2 % w/v) with different glycerol concentrations (10, 20, 30, 40, 50, 60, 75 % v/v).

SR-WAXS measurements were done by using the BL-40B2 spectrometer at the Japan Synchrotron Radiation Research Institute (JASRI, Harima, Japan) and by using the BL-10C spectrometer at the High Energy Accelerator Research Organization (KEK, Tsukuba, Japan). The X-ray wavelengths and the sample-to-detector distances were 51 cm for 0.75-Å X-ray and 4089 cm for 1.0-Å for X-ray at BL-40B2, and 190 cm for 1.49 -Å X-ray at BL-10C. The X-ray scattering intensity was recorded by the R-AXIS IV (30×30 cm^2^ in the area, 100-μm in pixel-resolution, from RIGAKU Co.) at both facilities. The exposure time was 10 seconds at BL-40B2, and 180 seconds at BL-10C. The temperature of the sample solutions contained in the sample cells was controlled in the temperature range from 25 °C to 85°C by using the temperature controller mK2000 of INSTEC co. While the measurements, the sample solutions were slowly oscillated to avoid some radiation damages. SANS measurement were carried out by using the D22 spectrometer at the research reactor of the Institut Laue-Langevin (ILL, Grenoble, France) and by using the BL15 TAIKAN spectrometer at the pulsed-neutron source of the Materials and Life Science Experimental Facility (MLF) at the Japan Proton Accelerator Research Complex (J-PARC, Tokai, Japan). The neutron wavelengths were 6 Å at ILL, and 0.5 - 6.0 Å at J-PARC. At both facilities, the sample solutions were contained in the quartz cells with 1 mm path length. The exposure time was around 10 – 30 minutes. Just before the scattering measurements, the sample solutions were filtered to remove some aggregates by using the centrifugal filter unit (Merck Co.) with the molecular-cut-off of 50 K Dalton.

The background correction for SANS data was done by an ordinary method [31]. WAXS data correction was done based on the method as reported previously [26, 27]. The radius of gyration *R*_g_ was determined by using the following equation [32].

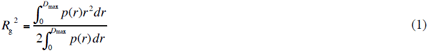

where the *p*(*r*) is the distance distribution function calculated by the Fourier inversion of the scattering curve

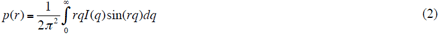

where *q* is the scattering vector (*q* = (4*π*/*λ*sin(*θ*/2); *θ*, the scattering angle; *λ*□□the X-ray wavelength. The maximum dimension *D*_max_ of the particle is determined from the *p*(*r*) function satisfying the condition *p*(*r*) = 0 for *r* > 0.

## 2. Results and discussion

### 2.1. Myoglobin structure depending on glycerol concentration observed by X-ray scattering

Figure 1 shows the WAXS curves of myoglobin depending on the glycerol concentration (0, 10, 20, 30, 40, 50, 60, 75 % v/v). As shown previously [26, 27], the WAXS curves of a protein in the different scattering regions mostly reflect the whole characteristics of the protein in the different hierarchical structure levels, that is, the quaternary and tertiary structures (*q* < ∼0.2 Å^-1^), the inter-domain correlation (∼0.25 Å^-1^< *q* < ∼0.5 Å^-1^), the intra-domain structure (∼0.5 Å^-1^ < *q* < ∼0.8 Å^-1^), and the secondary structure including the closely packed side chains (∼1.1 Å^-1^ < *q* < ∼1.9 Å^-1^), respectively. The scattering intensity of the particles in solutions depends on the difference between the scattering-density of the solute particle and that of the solvent [33]. It is so-called an excess average scattering-density, contrast (Δ*ρ*). In Figure 1, with the increase in the glycerol concentration, the scattering intensity in the small-*q* region decreases systematically due to the change of the contrast of myoglobin by the presence of glycerol. The change of the slope in the scattering curve below *q* = ∼0.2 Å^-1^ is not evidently seen, suggesting the preservation of the tertiary structure. On the other hand, the profile of the scattering curve in the *q* region from ∼0.25 Å^-1^ to ∼0.8 Å^-1^ shows a gradual change, which is evidently attributable to the change of the contrast, as shown in the following simulation. The profile of the scattering curve in ∼1.1 Å^-1^ < *q* < ∼1.9 Å^-1^ is mostly held. The above indicates both the intramolecular and secondary structures of myoglobin are almost preserved even by the existence of glycerol.

**Figure 1:**
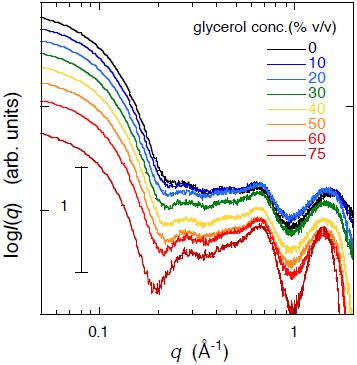
Wide-angle X-ray scattering (WAXS) curves of myoglobin depending on the glycerol concentration (0, 10, 20, 30, 40, 50, 60, 75 % v/v). The protein concentration is 2 % w/v in 10 mM HEPES buffer at pH 7.4 and 25 °C.

Figure 2 shows the distance distribution function, *p*(*r*), and the radius of gyration, *R*_*g*_, of myoglobin obtained by the Fourier transform of the scattering curve in Figure 1, where (A), *p*(*r*); (B), *R*_*g*_. The *p*(*r*) functions in Figure 2 (A) were calculated using Equation 1. The maximum diameter, *D*max, of the protein decreases from ∼54 Å at 0 % v/v glycerol to ∼47.5 Å at 50 % v/v glycerol and slightly increases to ∼48.5 Å at 75 % v/v. Whereas, the value of the peak-position, *p*(*r*)_max_, is mostly constant at ∼22 Å up to 50 % v/v glycerol, and turns to increasing to ∼25 Å at 75 % v/v glycerol. Figure 2 (B) shows the radius of gyration, *R*_*g*_. depending on the glycerol concentration, where the *R*_*g*_ values are obtained by using Equation 2. The *R*_*g*_ value once decreases from 16.9 ± 0.1 Å to 16.2 ± 0.1 Å at 50 % v/v glycerol, and increases to 16.9 ± 0.2 Å at 75 % v/v glycerol. The changing tendency of the *R*_*g*_ value agrees with that of the *D*max value in Figure 2 (A). The above WAXS results can be explained reasonably by the WAXS simulation based on the models of the preferential solvation and the preferential exclusion.

**Figure 2:**
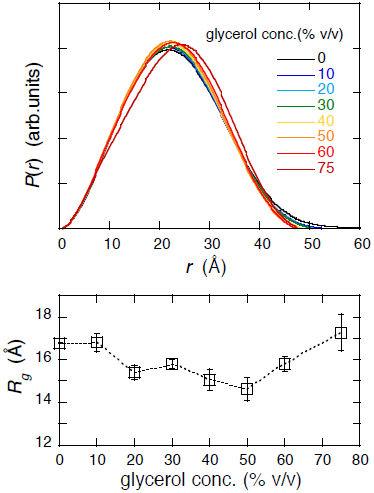
Distance distribution function, *p*(*r*), obtained by the Fourier transform of the scattering curve in Figure 1, and radius of gyration, *R*_*g*_, of myoglobin, where (A), *p*(*r*); (B), *R*_*g*_.

### 2.2 Simulation of wide-angle X-ray scattering (WAXS) curve and radius of gyration of myoglobin in crowder solution based on the preferential solvation model and the preferential exclusion model

As described above, the observed scattering curve of the solute particle depends on the difference between the scattering-density of the solute and that of the solvent, so-called contrast, Δρ. In the present experiment, the average scattering-density of the solvent varies depending on the glycerol concentration. To execute the WAXS simulation, we should estimate the variation of the Δρ in glycerol-water mixed solutions. Therefore, we have measured the mass-densities of the glycerol solvents to determine those scattering densities. Figure 3 (A) shows the observed densities of the water solvents containing glycerol at different concentrations (% v/v) by using the electric balance (Sartorius R200D). The mass density shows a good linearity in the concentration. The average scattering-densities (electron densities) of proteins is in the range of ∼11.7 - ∼12.0 ×10^10^ cm^-2^ (∼0.416 eÅ^-3^ – 0.427 eÅ^-3^) for X-ray [33]. Based on the crystal structure of myoglobin (code no. 1WLA registered in PDB [34]), the average scattering-density of myoglobin can be calculated to be 11.9 ×10^10^ cm^-2^ (0.424 eÅ^-3^) for X-ray. Figure 3 (B) shows the glycerol-concentration dependence of the contrast of myoglobin for X-ray both in cm^-2^ and eÅ^-3^ units.

**Figure 3:**
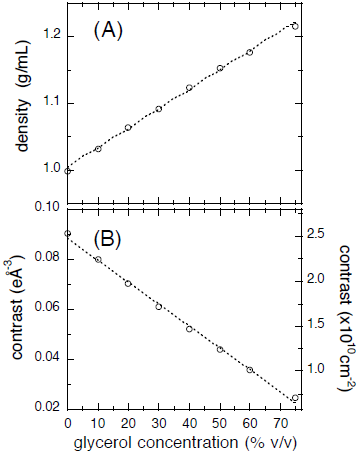
Observed mass density of the water solvent containing glycerol at different concentrations (% v/v) and estimated value of the excess average scattering-density, so-called contrast for X-ray, of myoglobin in glycerol solution, where (A), mass density; (B), contrast.

As shown in Figure 3 (B), the presence of glycerol in the solvent changes the contrast of the protein and results in the change of the scattering curve. By the comparison of the theoretical calculation of the scattering curve with the experimental one, we can discuss the detail of the effect of glycerol on the protein structure observed experimentally. The theoretical WAXS simulations were executed by using the CRYSOL program [35]. The CRYSOL program is based on the spherical-harmonics expansion method [33], and takes into account of the existence of the hydration shell by using the atomic coordinates of a protein in the PDB. In the CRYSOL program, the contrast of the hydration shell is a variable, and its width is set to be 3 Å as a default value. This program is known to reproduce experimental X-ray scattering curves of proteins in solutions very well [36]. The default value of the hydration-shell width would be reasonable. As shown by the previous studies on protein hydration [37], the amount of protein-hydration corresponds to that of water molecule covering protein surface with at least two layers of water (so-called, strongly bound water layer and weakly bound water layer). Here we used the PDB file with the number of 1WLA of myoglobin for the CRYSOL calculation. The present theoretical simulation was performed assuming the following two different cases that can happen on the protein structure and its hydration-shell caused by glycerol. The first case is that the preferential replacement of hydration-shell water molecules with glycerol molecules occurs (preferential solvation effect model [38]). In this case, the preferential arrangement of glycerol molecules surrounding the protein surface causes the increase in the scattering density of the solvation-shell (hydration-shell) of the protein according to the rise of the glycerol concentration. The second case is that glycerol molecules are preferentially excluded from the hydration-shell region of the protein due to the hydration repulsion force. In this case, the hydration-shell density is preserved and keeps a constant value in spite of the rise of glycerol concentration. The above two cases correspond to two extreme ones that would occur physicochemically induced by co-solutes. In other words, the former case and the later one are equivalent to the assumptions of the partial replacement of hydrated water with glycerol and the non-penetration of glycerol into the hydration shell, respectively.

Figure 4 shows the simulated WAXS curves with the rise of the average scattering density (electron density) of the solvent which is proportional to the glycerol concentration, where (A), preferential solvation model; (B), preferential exclusion model. The variable range of the average electron density of the solvent in Figure 4 is compatible with that of the glycerol concentration from 0 to 75 % v/v. Based on two models, in Figure 4 (A), the average electron density of the hydration-shell region was set to be 1.1 times larger than that of the glycerol solvent; in Figure 4 (B), the average electron density of the hydration-shell region was set to be constant, namely, 1.1 times larger than that of free water. Compared with the experimental scattering curve in the range form *q* = ∼0.2 Å^-1^ to *q* = ∼0.3 Å^-1^, it can be seen that the tendency of the change is more similar to that of the simulated WAXS curve in Figure 4 (B) than in Figure 4 (A). In addition, the change of the small-angle scattering intensity (*q* < 0.1 Å^-1^) in Figure 4 (B) with the rise of the solvent electron density is much larger than in Figure 4 (A).

**Figure 4:**
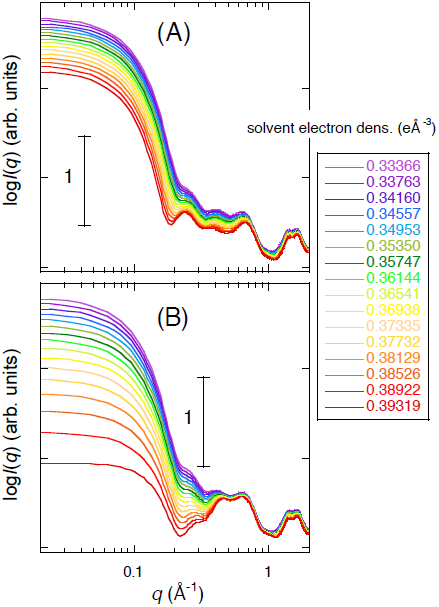
Simulated WAXS curve depending on the average scattering density (electron density) of the solvent that corresponds to the rise of the glycerol concentration. The variable range of the average electron density of the solvent is compatible with that of the glycerol concentration from 0 to 75 % v/v in Figure 1. The simulation using the CRYSOL program was done based on two different models. Namely, (A), preferential solvation effect model (preferential replacement of hydration-shell water molecules of the protein surface with glycerol molecules); (B), preferential exclusion effect model (preferentially exclusion of glycerol molecules from the hydration-shell region of the protein).

Figure 5 shows the normalized values of the experimental and theoretical square-root of the zero-angle scattering intensity, *I*(0)^1/2^ and *R*_*g*_, where (A), *I*(0)^1/2^; (B), *R*_*g*_. The *I*(0)^1/2^ value is well known to be proportional to the product of the contrast and the volume of the solute particle [26]. In Figure 5 (A), the difference between the slopes of two linear relationships in the theoretical *I*(0)^1/2^ values indicates that the *I*(0)^1/2^ value in the preferential exclusion model decreases much significantly compared with that in the preferential solvation model. Below the glycerol concentration of ∼40 % v/v, The decreasing tendency of the experimental *I*(0)^1/2^ and *R*_*g*_ with the rise of the glycerol concentration is mostly quantitatively reproduced by the preferential exclusion model, as shown in Figure 5. On the other hand, above the glycerol concentration of ∼50 % v/v the deviation of both experimental values from the theoretical ones based on the preferential exclusion model become to be evidently seen. It should be noted that the theoretical *R*_*g*_ values based on two different models clearly show opposite tendencies on the rise of the glycerol concentration. The deviation of the experimental *I*(0)^1/2^ and the increasing tendency of the experimental *R*_*g*_ value above ∼50 % v/v can be explained by a partial replacement or penetration of the protein hydration-shell water molecules with glycerol molecules, namely by the preferential solvation model.

**Figure 5:**
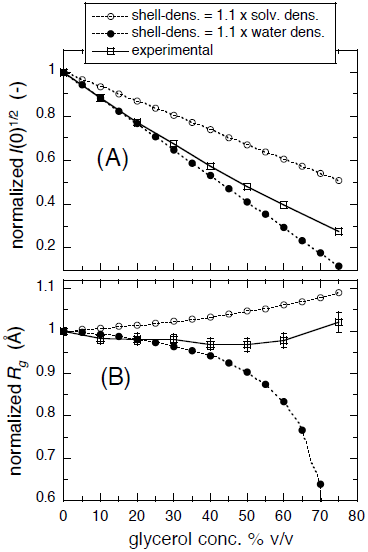
Normalized values of the experimental and theoretical square-root of the zero-angle scattering intensity, *I*(0)^1/2^ and *R*_*g*_, where (A), *I*(0)^1/2^; (B), *R*_*g*_.

The above experimental and simulation results suggest that glycerol molecules are preferentially excluded from the protein surface region up to the concentration of ∼40 % v/v and that at the concentration over ∼50 % v/v glycerol molecules partially penetrate into the hydration region of the protein. The above interpretation is strongly supported by the following experimental and simulation results of neutron scattering.

### 2.3. Effect of glycerol on protein structure observed by neutron scattering

In the case of neutron scattering, various types of the contrast-variation method are available to determine the structure of a particle in solution [39]. We have employed the inverse-contrast variation method to observe the effect of glycerol on the protein structure. This method using deuterated materials can avoid or minimize the artificial effect on the scattering curves caused by the addition of co-solute molecules. The mixture of non-deuterated glycerol (h-glycerol) and deuterated glycerol (d-glycerol, 98 atom % D) was used. As we know the molecular volume of glycerol from the mass-density measurement, the average-scattering densities of h-glycerol and 98 % d-glycerol can be calculated. The average scattering-density of the mixture of h-glycerol / 98 % d-glycerol = 21.25 / 78.75 (v/v) corresponds to that of 100 % D_2_O. This mixture can minimize both the coherent and incoherent background scatterings from glycerol molecules and hydrogen atoms. In addition, by the use of this mixture, the increase of the glycerol content does not change the contrast of the protein. While, in the case of X-ray scattering, the change of the contrast is unavoidable, as shown in the above section. In other words, we are able to observe selectively only the protein structure including its hydration shell under the above experimental condition.

Figure 6 shows the glycerol-concentration dependence of the neutron scattering curve of myoglobin, where the insert is the distance distribution function, *p*(*r*). The glycerol-concentration was varied from 0 % v/v to 60 % v/v. The maximum diameter, *D*_max_, in the *p*(*r*) function mostly holds in the range from 49 Å to ∼50 Å. Figure 7 shows the square-root of the zero-angle scattering intensity, *I*(0)^1/2^, and the *R*_*g*_ value obtained from Figure 6, where (A), *I*(0)^1/2^; (B), *R*_*g*_. Although both changes in the scattering curve and in the *p*(*r*) function are minor, the *I*(0)^1/2^ value in Figure 7 (A) shows a slightly increasing tendency in ∼4 % at 60 % v/v. The *R*_*g*_ value in Figure 7 (B) also shows a slight increase from 13.4 ± 0.1 Å to 13.6 ± 0.1 Å. These changes are reasonably explained by the change in the hydration-shell density as shown below. It should be noted that the difference between the absolute *R*_*g*_ values obtained by SANS and SR-WAXS are essentially attributable to the difference in the contrast profile of myoglobin in D_2_O (for SANS) and H_2_O (for SR-WAXS).

**Figure 6:**
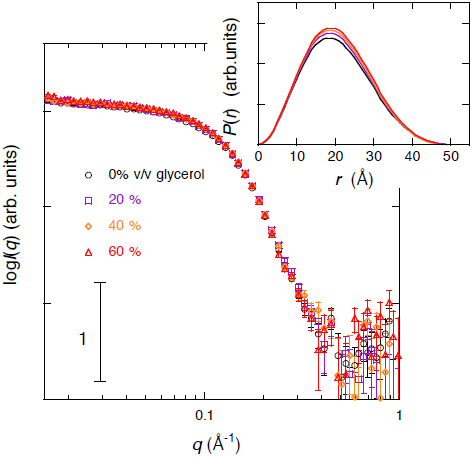
Neutron scattering curve depending on the glycerol concentration. The protein concentration is 2 % w/v in 100 % D_2_O, 10 mM Hepes, at pH 7.4 and 25 °C. Glycerol is the mixture of [h-glycerol]/[98 % d-glycerol] = 21.25/78.75. The average scattering density of the glycerol mixture matches with that of 100 % D_2_O. The insert shows the distance distribution function.

**Figure 7:**
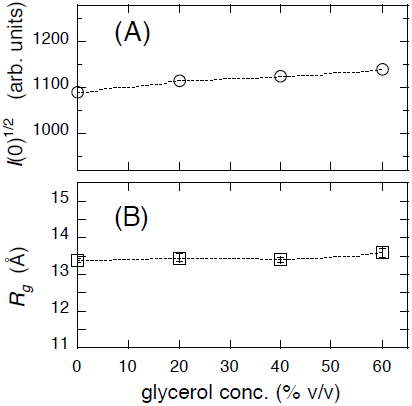
Square-root of the zero-angle scattering intensity, *I*(0)^1/2^, and radius of gyration, *R*_*g*_, obtained from Figure 6, where (A), *I*(0)^1/2^; (B), *R*_*g*_.

### 2.4. Fitting by theoretical neutron scattering curve and estimation of hydration-shell density

In the present neutron scattering experiment, we have applied the inverse-contrast variation method to avoid the artifact of the change of the contrast caused by the addition of glycerol. Namely, even by the rise of the glycerol concentration, both the average scattering density of the solvent and the contrast of the protein did not change. Therefore, we can effectively use the theoretical fitting procedure to reproduce the experimental scattering curve by the CRYSON program. The CRYSON program is a reimplementation of CRYSOL, adapted to work with small-angle neutron scattering data. Figure 8 shows the optimized simulated neutron scattering curve with the experimental one at each glycerol concentration. The discrepancy between theoretical and experimental curves defined by *χ*^2^ value was in the range from 0.92 to 1.4. The estimated contrast of the hydration (solvation) shell by the CRYSON fitting is shown in Figure 9. The hydration-shell contrast preserves the value of 0.61 ×10^12^ cm^-2^ (corresponding to 9.5 % higher than the average scattering-density of D_2_O) up to 40 % v/v glycerol and decreases to 0.48 ×10^12^ cm^-2^ at 60 % v/v glycerol. Under the present experimental condition, the average scattering density of glycerol matches with that of D_2_O. Therefore, the replacement of the hydration-shell water surrounding the protein surface with glycerol reduces the hydration-shell contrast. The decrease of the hydration-shell contrast directly suggests the preferential replacement of hydration-shell water molecules with glycerol molecules. The fraction of the replacement of the hydration-shell water with glycerol would be around 20 %.

**Figure 8:**
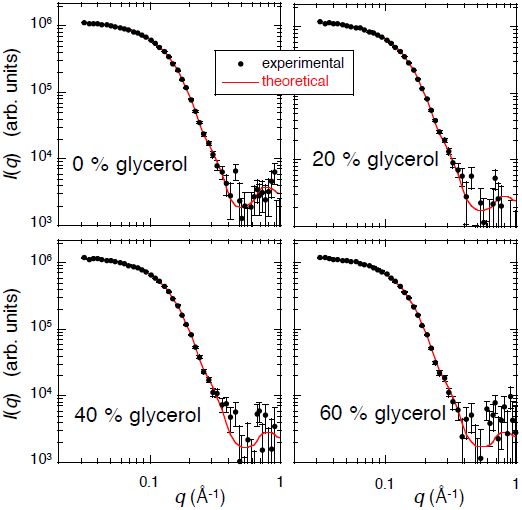
Theoretical scattering curve to fit experimental one in Figure 7 by using CRYSON program. The solid lines and the marks correspond to the optimized simulated neutron scattering curves and the experimental data, respectively. The discrepancy defined by *χ*^2^ value is in the range from 0.92 to 1.4.

**Figure 9:**
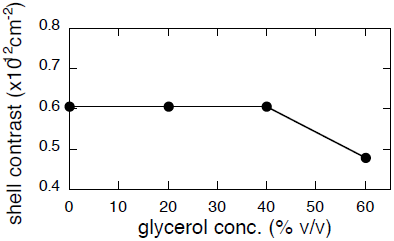
Estimated contrast of the hydration (solvation) shell obtained by the CRYSON fitting shown in Figure 8.

The effect of the change in the hydration-shell contrast on the scattering curve was estimated by the scattering function simulation using the CRYSON program. In Figure 10, the hydration-shell contrast varied from 0.064 ×10^12^ cm^-2^ to 0.04 ×10^12^ cm^-2^. When the hydration-shell contrast changes from 0.61 ×10^12^ cm^-2^ to 0.48 ×10^12^ cm^-2^, the *I*(0)^1/2^ and *R*_*g*_ values increase in ∼3 % and in ∼2 %, respectively. These theoretical increments mostly agree with those shown in Figure 7.

**Figure 10:**
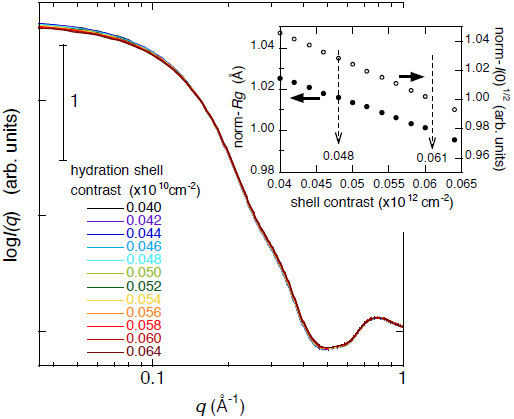
Theoretical neutron scattering curve depending on the hydration-shell contrast obtained by the CRYSON program. The hydration-shell contrast varied from 0.064 ×10^12^ cm^-2^ to 0.04 ×10^12^ cm^—2^. The insert shows the *I*(0)^1/2^ and *R*_*g*_ values depending on the hydration-shell contrast. For the change of the hydration-shell contrast from 0.61 ×10^12^ cm^-2^ to 0.48 ×10^12^ cm^—2^ (as shown in Figure 9), the *I*(0)^1/2^ and *R*_*g*_ values increase in ∼3 % and in ∼2 %, respectively.

Thus, the result of neutron scattering suggests that at below 40 % v/v glycerol the preferential exclusion of glycerol molecules from the hydration region on the protein surface is dominant and that at higher concentration the preferential solvation (penetration) of glycerol molecules becomes to occur locally. This result agrees well with that obtained from the WAXS experiment and theoretical simulation in the above section.

### 2.5. Thermal structure stability and helix-to-sheet transition in glycerol solution

Figure 11 shows the temperature dependence of WAXS curve of myoglobin, where A, in the non-crowding solvent; B, in 20 % v/v glycerol solvent. As already mentioned in the section of 2.1, the observed WAXS scattering data covers the distinct levels of the protein structure from the tertiary structure to the secondary one. The scattering data at *q* < ∼0.2 Å^-1^, at ∼0.25 Å^-1^ < *q* < ∼0.5 Å^-1^, at ∼0.5 Å^-1^ < *q* < ∼0.8 Å^-1^, and at ∼1.1 Å^-1^ < *q* < ∼1.9 Å^-1^ reflect the tertiary structure, the inter-domain correlation, the intra-domain structure, and the secondary structure, respectively. Therefore, we can analyze the structural transition feature at different hierarchical levels separately. The shoulder at *q* = ∼0.58 Å^-1^ and the peak at *q* = ∼1.34 Å^-1^ become to be evident with the elevation of temperature above ∼75 °C. As reported previously [24, 25, 40], the former and the latter are typical phenomena in the initial process of amyloid transition, namely, the formations of the pleated sheet stacking and the appearance of cross-β structure (helix-to-sheet transition), respectively. The appearance of the shoulder at *q* = ∼0.58 Å^-1^ and the peak at *q* = ∼1.34 Å^-1^ are as same as that observed in the amyloid transition of apomyoglobin [28, 29]. The *q* values of the positions of the shoulder and the peak correspond to ∼10.8 Å and ∼4.69 Å in the real space distance, respectively. These values agree with those reported previously within experimental errors [24]. Figure 12 shows the temperature dependence of the distance distribution function, *p*(r), obtained by applying Eq. 1 to the scattering curves in Figure 11, where A and B are as in Figure 11. At low temperature, in the short-distance region, the *p*(r) function shows the bell-shape profile having the peak at ∼20 Å and the first minimum at ∼45 Å, which reflects the gradual change of the monomer structure of myoglobin. In the long-distance region, above a certain temperature, the *p*(r) function becomes to show another modulated hump, or shoulder, suggesting the appearance of oligomeric aggregates, namely, amyloid-like aggregates. As is clearly shown in Figure 12, the presence of glycerol suppressed the thermal unfolding and oligomerization of the protein.

**Figure 11:**
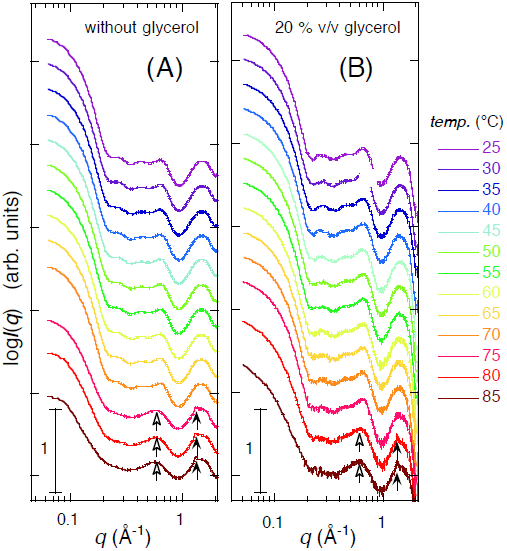
Temperature dependence of the WAXS curve of myoglobin (2 % w/v in 10 mM HEPES buffer at pH 7.4) in the non-crowding solvent, A, and in 20 % v/v glycerol solvent, B. The fill and open arrows indicate the typical features in the scattering curve that appear in the initial process of amyloid transition. The shoulder at *q* = ∼0.58 Å^-1^ and peak at *q* = ∼1.34 Å^-1^ correspond to the formations of the pleated sheet stacking and the appearance of cross-β structure (helix-to-sheet transition), respectively.

**Figure 12:**
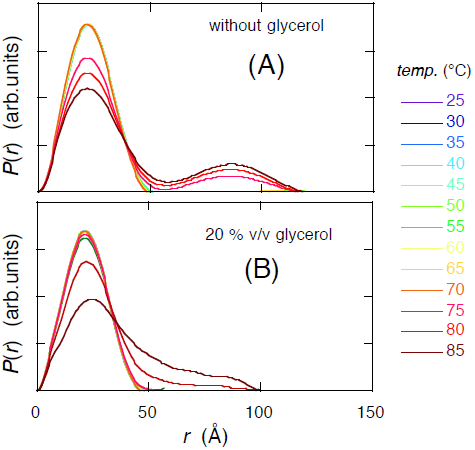
Temperature dependence of the distance distribution function of myoglobin calculated from the WAXS curves in Figure 11. A and B are as in Figure 11.

The observed WAXS scattering data covers the distinct levels of the protein structure from the tertiary structure to the secondary one. Namely, as shown previously [26, 27], the scattering data at *q* < ∼0.2 Å^-1^, at ∼0.25 Å^-1^ < *q* < ∼0.5 Å^-1^, at ∼0.5 Å^-1^ < *q* < ∼0.8 Å^-1^, and at ∼1.1 Å^-1^ < *q* < ∼1.9 Å^-1^ mostly reflect the tertiary structure, the inter-domain correlation, the intra-domain structure, and the secondary structure including the closely packed side chains, respectively. Therefore, we can analyze the transition feature at different hierarchical structural levels separately.

The transition-multiplicity analysis (TMA) [27] is applicable to characterize the thermal stability in the different hierarchical-structure levels. The TMA method is based on the principle that the scattering curves in different *q*-regions correspond to the structures of an object in the different hierarchical-structure levels. The structural transitions in the different hierarchical-structure levels do not necessary proceed cooperatively. By using Equation 2 we can analyze the transition cooperativity among the different hierarchical-structure levels [23, 41, 42].

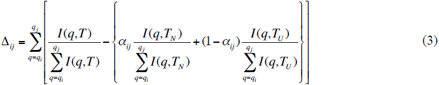

Where *I*(*q*,*T*_*N*_),*I*(*q*,*T*_*U*_) and *I*(*q*,*T*) are the scattering profiles at the initial, final, and intermediate temperatures in a defined *q*-range of *q*_i_ *q*_j_ Å^-1^, respectively. The scattering curve *I*(*q*,*T*) in a defined *q*-range at an intermediate temperature is fitted by using a linear combination of *α*_*ij*_ *I* (*q*, *T*_*N*_) and (1-*α*_*ij*_)*I*(*q*,*T*_*U*_) at the initial and final temperatures. The factor *αij* is determined by minimizing theΔ_*ij*_ value in Equation 2. Thus, the *α*_*ij*_ value corresponds to the molar fraction of the initial temperature state at the intermediate temperature *T* in the distinct hierarchical structure level. In the TMA results in Figure 13, the selected *q*-ranges are 0.05 – 0.1 Å^-1^, 0.25 – 0.8 Å^-1^, and 1.2 – 1.8 Å^-1^ that correspond to the tertiary structure, the inter-domain correlation and the intra-domain structure, and the secondary structure, respectively. The midpoint temperature of the thermal transition, *T*_m_, can be determined by the temperature at *α* = 0.5 in Figure 13. In the case of myoglopin in the solvent without glycerol, the thermal structural transition proceeds cooperatively among all different hierarchal structural levels at *T*_m_ = ∼75 °C. Whereas, in the glycerol solvent, the *T*_m_ value rises, suggesting that the transition cooperativity between the tertiary structure and the internal and secondary structures is weakened. The *T*_m_ value is 77.6 °C for the internal and secondary structure levels, and 79.5 °C for the tertiary structure level. The difference in the *T*_m_ values suggests that the presence of glycerol in the solvent stabilizes the tertiary structure much more than the internal and secondary structures.

**Figure 13:**
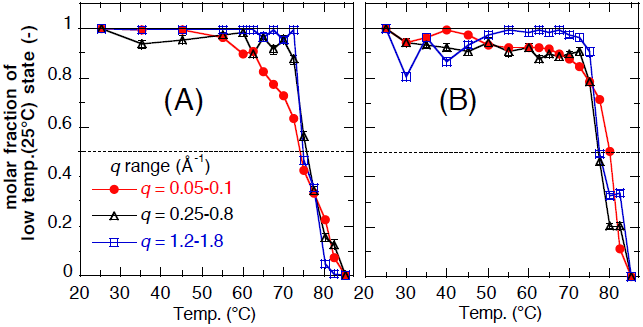
Molar fraction *α* (*α* ≤ 1) of the initial structure (at 25 °C) of myoglobin at an intermediate temperature in the heating process determined from the scattering curves in different *q*-ranges by using the TMA analysis, where A, in the non-crowding solvent; B, in 20 % v/v glycerol solvent. The marks of circle, triangle, and square show the *α* values using the *q*-ranges of 0.05 – 0.1 Å^-1^, 0.2 – 0.8 Å^-1^, and 1.2 – 1.8 Å^-1^, respectively. These ranges correspond to the different hierarchal structure levels of myoglobin, namely, the tertiary structure, the inter-domain correlation and the intra-domain structure, and the secondary structure including the closely packed side chains, respectively. The dashed line indicates *α* = 0.5. The intersection temperature between the dashed line and the *α* value is the transition-midpoint temperature for each hierarchical structure.

## 3. Conclusion

By the complementary use of SR-WAXS and SANS methods and by the theoretical simulation of scattering functions, we have clarified the effect of glycerol molecules on the myoglobin structure, its hydration layer and thermal stability as follows. At the concentration of glycerol lower than ∼40 % v/v, the WAXS and theoretical simulation results suggest that glycerol molecules are preferentially excluded from the hydration-shell region of the protein (preferential exclusion) to hold the hydration-shell. This was confirmed directly as the preservation of the hydration-shell density by the SANS results using the inverse-contrast variation method and by the theoretical fitting with those experimental data. Due to the preservation of the hydration-shell by glycerol, the protein structure is stabilized against temperature elevation, especially for the tertiary structure of myoglobin compared with that for the internal and secondary structures. At the concentration of glycerol higher than ∼50 % v/v, the partial penetration or replacement of glycerol molecules into or with the hydrated water molecules surrounding the protein surface (preferential solvation). The present results clearly show that the stabilizing effect of the protein structure by glycerol is caused by the protective action of the hydration shell of the protein.

In this study we have shown the small-molecular-crowing effect on the protein structure and stability. Although there are many previous studies of the crowding effect on biological function such as enzymatic synthesis, signal transduction, and so on, using glycerol as a crowder agent, the mechanism of macromolecular crowding effect, especially in living cell should be more complicated by the diversity of molecular species and structural properties [43]. Therefore, further development and combination of experimental and theoretical methods treating macromolecular crowding environment are necessary. The combination of SR-WASX method and the inverse-contrast method of neutron scattering might offer certain experimental advantages for studying macromolecular crowding effect on protein structures.

## Notes

The authors declare no competing financial interest.

## ACKNOWLEDGMENTS

The X-ray scattering experiments were performed under the approval of the program advisory committee of the Japan Synchrotron Radiation Research Institute (Proposal No. 2016A1487 & 2017B1435) and under the approval of the PF program advisory committee (Proposal No. 2015G518). The neutron scattering experiment was done under the approval of the Neutron Scattering Program Advisory Committee (Proposal No. 2014P0501) and under the approval of the ILL Program Advisory Committee (Proposal No. 8-03-740). This research project was partly supported by the Grants-in-Aid for Scientific Research of JSPS (the Japan Society of the Promotion of Science) (Proposal No. 16K13722).

## References

1. Gekko, K., and S. N. Timasheff. 1981. Mechanism of protein stabilization by glycerol: Preferential hydration in glycerol-water mixtures. Biochemistry 20:4667–4676.

2. Calhoun, D. B., and S. W. Englander. 1985. Internal protein motions, concentrated glycerol, and hydrogen exchange studied in myoglobin. Biochemistry 24:2095–2100.

3. Farnum, M. and C. Zukoski. 1999. Effect of Glycerol on the Interactions and Solubility of Bovine Pancreatic Trypsin Inhibitor. Biophys. J. 76:2716–2726.

4. Vagenende, V., Yap, M. G. S., and B. L. Trout. 2009. Mechanisms of Protein Stabilization and Prevention of Protein Aggregation by Glycerol. Biochemistry 48: 11084–11096.

5. Caliskan, G., Mechtani, D., Roh, J. H., Kisliuk, A., Sokolov, A. P., Azzam, S., Cicerone, M. T., Lin-Gibson, S., and I. Peral. 2004. Protein and solvent dynamics: How strongly are they coupled? J. Chem. Phys. 121:1978–1983.

6. Ronsina, O., Caroli, C., and T. Baumberger. 2017. Preferential hydration fully controls the renaturation dynamics of collagen in water-glycerol solvents. Eur. Phys. J. E 40:55, DOI10.1140/epje/i2017-11545-1.

7. Pain, R. H. ed.; Mechanisms of protein folding; Oxford University Press, 2000.

8. Dobson, C. M. 2003. Protein folding and misfolding, Nature 426:884–890.

9. Goodsell, D. S. 1991. Inside a living cell, Trend Biochem. Sci. 16:203–206.

10. Davis-Searles, P. R., and A. J. Saunders. 2001. Interpreting the effect of small unchaged solutions on protein-folding equilibria. Annu. Rev. Biophys. Biomol. Struct. 30: 271–306.

11. Rosgen, J., Pettitt, B. M., and D. W. Bolen. 2005. Protein Folding, Stability, and Solvation Structure in Osmolyte Solutions. Biophys. J. 89:2988–2997.

12. Rosgen, J., Pettitt, B. M., and D. W. Bolen. 2007. An analysis of the molecular origin of osmolyte-dependent protein stability, An analysis of the molecular origin of osmolyte-dependent protein stability. Protein Sci. 16:733–743.

13. Fayer, M. D. 2012. Dynamics of water interacting with interfaces, molecules, and ions. Accounts of Chemical Reseach 45: 3–14.

14. Makarov, V. A., Andrews, B. K., Smith, P. E., and B. M. Pettitt. 2000. Residence Times of Water Molecules in the Hydration Sites of Myoglobin. Biophys. J. 79:2966– 2974.

15. Zaccai, G. 2000. How soft is a protein? A protein dynamics force constant measured by neutron scattering. Science 288:1604–1607.

16. Gabel, F., Bicout, D., Lehnert, U., Tehei, M., Weik, M., and G. Zaccai. 2002. Protein dynamics studied by neutron scattering. Q. Rev. Biophys. 35:327–367.

17. Tehei, M., Franzetti, B., Wood, K., Gabel, F., Fabiani, E., Jasnin, M., Zamponi, M., Oesterhelt, D., and G. Zaccai. 2007. Ginzburg, M. Neutron scattering reveals extremely slow cell water in a Dead Sea organism. Proc. Nat. Acad., Sci., USA 104:766–771.

18. Froelich, A., Gabel, F., Jasnin, M., Lehnert, U., Oesterhelt, D., Stadler, A. M., Tehei, M., Weik, M., Wood, K., and G. Zaccai. 2009. From shell to cell: neutron scattering studies of biological water dynamics and coupling to activity. FARADAY DISCUSSIONS 141:117–130.

19. Sukenik, S., Sapir, L., and D. Harries. 2013. Balance of enthalpy and entropy in depletion forces. Current Opinion in Colloid & Interface Science 18:495–501.

20. Dastidar, S. G., and C. Mukhopadhyay. 2003. Structure, dynamics, and energetics of water at the surface of a small globular protein: A molecular dynamics simulation. Phys. Rev. E 68:21921–21930.

21. Harada, R., Sugita, Y., and M. Feig. 2012. Protein Crowding Affects Hydration Structure and Dynamics. J. Am. Chem. Soc. 134:4842–4849.

22. Ohno, Y., Yokota, R., Koyama, H., Morimoto, G., Hasegawa, A., Masumoto, G., Okimoto, N., Hirano, Y., Ibeid, H., Narumi, T., and M. Taiji. 2014. Petascale molecular dynamics simulation using the fast multipole method on K computer. Comp. Phys. Comm. 185:2575–2585.

23. Wang, P., Yu, I., Feig, M., and Y. Sugita. 2017. Influence of protein crowder size on hydration structure and dynamics in macromolecular crowding. Chemical Physics Letters 671:63–70.

24. Fändrich, M., Fletcher, M. A., and C. M. Dobson. 2001. Amyloid fibrils from muscle myoglobin. Nature 410:165–166.

25. Fändrich, M., Forge, V., Buder, K., Kittler, M., Dobson, C. M., and S. Diekmann. 2003. Myoglobin forms amyloid fibrils by association of unfolded polypeptide segments. Proc. Nat. Acad., Sci., USA 100:15463–15468.

26. Hirai, M., Iwase, H., Hayakawa, T., Miura, K., and K. Inoue. 2002. Structural hierarchy of several proteins observed by wide-angle solution scattering. J. Synchrotron Rad. 9:202–205.

27. Hirai, M., Koizumi, M., Hayakawa, T., Takahashi, H., Abe, S., Hirai, H., Miura, K., and K. Inoue. 2004. Hierarchical map of protein unfolding and refolding at thermal equilibrium revealed by wide-angle X-ray scattering. Biochemistry 43:9036–9049.

28. Onai, T., Koizumi, M., Lu, H., Inoue, K., and M. Hirai. 2007. Initial process of amyloid formation of apomyoglobin and effect of glycosphingolipid GM1. J. Appl. Cryst. 40:s184–s189.

29. Fabian, E., Stadler, A. M., Madern, D., Koza, M. M., Tehei, M., Hirai, M., and G. Zaccai. 2009. Dynamics of apomyoglobin in the transition and of partially unfolded aggregated protein. Eur. Biophys. J. 38:237–244.

30. Knoll, W., Schmidt, G., and K. Ibel. 1985. The inverse contrast variation in smallangle neutron-scattering. J. Appl. Cryst. 18:65–70.

31. Hirai, M., Iwase, H., Hayakawa, T., Koizumi, M., and H. Takahashi. 2003. Determination of Asymmetric Structure of Ganglioside-DPPC Mixed Vesicle Using SANS, SAXS and DLS. Biophys. J. 85:1600–1610.

32. Glatter, O. in Small-Angle X-ray Scattering (Glatter, O., and Kratky, O., Eds.) pp 119–196, Academic Press, London, 1982.

33. Stuhrmann, H. B. 1978. Miller, A. Small-angle scattering of biological structures. J. Appl. Cryst. 11:325–345.

34. Maurus, R., Overall, C. M., Bogumil, R., You, Y., Mauk, A. G., Smith, M., and G. D. Brayer. 1997. A Myoglobin Variant with a Polar Substitution in a Conserved Hydrophobic Cluster in the Heme Binding Pocket, Biochem. Biophys. Acta 134:1–13.

35. Svergun, D. I., Barberato, C., and M. H. J. Koch. 1995. CRYSOL-a program to evaluate X-ray solution scattering of biological macromolecules from atomic coordinates, J. Appl. Cryst. 28:768–773.

36. Svergun, D. I., Richard, S., Koch, M. H. J., Sayers, S., Kuprin, S., and G. Zaccai. 1998. Protein hydration in solution: Experimental observation by X-ray and neutron scattering, Proc. Natl. Acad. Sci. USA 95:2267–2272.

37. Rupley, J. A., and G. Careri. 1991. Protein hydration and function. Adv. Protein Chem. 41:37–172.

38. Auton, M., Bolen, D. W., and J. Rosgen. 2008. Structural thermodynamics of protein preferential solvation. Protein-Structure Function and Bioinformatics 73:802–813.

39. Hirai, M. ‘Contrast Variation’ in Neutrons in Soft Matter, pp. 351-382, edited by T. Imae, T. Kanaya, M. Furusaka, and N. Torikai, John Wiley & Sons, Inc., 2011.

40. Malinchik, S. B., Inouye, H., Szumowski, K. E., and D. A. Kirschner. 1998. Structural analysis of Alzheimer’s beta(1-40) amyloid: Protofilament assembly of tubular fibrils. Biophys. J. 74:537–545.

41. Hirai, M., Arai, S., and H. Iwase. 1999. Complementary analysis of thermal transition multiplicity of hen egg-white lysozyme at low pH using X-ray scattering and scanning calorimetry. J. Phys. Chem. B 103:549–556.

42. Hirai, M., Sato, S., Kimura, R., Hagiwara, Y., Kawai-Hirai, R., Ohta, N., Igarashi, N., and N. Shimizu. 2015. Effect of protein-encapsulation on thermal structural stability of liposome composed of glycosphingolipid/cholesterol/phospholipid. J. Phys. Chem. B 119:3398–3406.

43. Gnutt, D., Gao, M., Brylski, O., Heyden, M., and S. Ebbinghaus. 2015. Excluded-volume effects in living cells. Angew. Chem. Int. Ed. 54:2548 –2551.

